# A D-2-Hydroxyglutarate dehydrogenase mutant reveals a critical role for ketone body metabolism in *Caenorhabditis elegans* development

**DOI:** 10.1101/2022.05.16.492161

**Authors:** Olga Ponomarova, Hefei Zhang, Xuhang Li, Shivani Nanda, Thomas B. Leland, Bennett W. Fox, Gabrielle E. Giese, Frank C. Schroeder, L. Safak Yilmaz, Albertha J.M. Walhout

## Abstract

In humans, mutations in D-2-hydroxyglutarate (D-2HG) dehydrogenase (D2HGDH) result in D-2HG accumulation, delayed development, seizures, and ataxia. While the mechanisms of 2HG-associated diseases have been studied extensively, the endogenous metabolism of D-2HG remains unclear in any organism. Here, we find that, in *Caenorhabditis elegans*, D-2HG is produced in the propionate shunt, which is transcriptionally activated when flux through the canonical, vitamin B12-dependent propionate breakdown pathway is perturbed. Deletion of the D2HGDH ortholog, *dhgd-1*, results in embryonic lethality, mitochondrial defects, and the upregulation of ketone body metabolism genes. Viability can be rescued by RNAi of *hphd-1*, which encodes the enzyme that produces D-2HG, or by supplementing either vitamin B12 or the ketone body 3-hydroxybutyrate (3HB). Altogether, our findings support a model in which *C. elegans* relies on ketone bodies for energy when vitamin B12 levels are low, and in which a loss of *dhgd-1* causes lethality by limiting ketone body production.

**HIGHLIGHTS:**

- D-2-hydroxyglutarate is produced by HPHD-1 in the propionate shunt pathway
- DHGD-1 recycles 2-hydroxyglutarate to sustain flux through the propionate shunt
- *dhgd-1* loss perturbs ketone body metabolism and causes embryonic lethality
- 3-Hydroxybutyrate, vitamin B12 or *hphd-1* RNAi rescue *dhgd-1* mutant lethality

## INTRODUCTION

The metabolite 2HG occurs as two enantiomers, L-2HG and D-2HG, each of which is oxidized by a specific dehydrogenase. Mutations in these dehydrogenases result in the inborn errors of human metabolism, L- and D-2-hydroxyglutaric aciduria, respectively (Faiyaz-Ul-Haque et al., 2014; Veiga-da-Cunha et al., 2020). These diseases cause the accumulation of 2HG in bodily fluids, delayed development, neurological and muscle dysfunction, and early death (Kranendijk et al., 2012). Both enantiomers are oncometabolites, but they are produced differently and have distinct effects on metabolism and physiology. L-2HG accumulates during hypoxia and is produced by malate and lactate dehydrogenases (Intlekofer et al., 2015; Intlekofer et al., 2017). Both L-2HG and D-2HG accumulate in humans with mutated mitochondrial citrate transporter (Pop et al., 2018), and an underlying mechanism of this disorder was proposed by studies in model organism *Drosophila* (Li et al., 2018). D-2HG drives oncogenic transformation in cells with neomorphic mutations in the isocitrate dehydrogenases IDH1 and IDH2 (Chen et al., 2013; Dang et al., 2009). D-2HG inhibits multiple enzymes, including alpha-ketoglutarate (αKG) dependent dioxygenases (Chowdhury et al., 2011; Xu et al., 2011), BCAT transaminases (McBrayer et al., 2018), αKG dehydrogenase (Karlstaedt et al., 2016), and ATP synthase (Fu et al., 2015). D-2HG can be produced by several enzymes, including the hydroxyacid-oxoacid transhydrogenase ADHFE1 (Struys et al., 2005), the phosphoglycerate dehydrogenase PHGDH (Fan et al., 2015) and wild type IDH1 and IDH2 (Intlekofer et al., 2017). However, it remains unclear if D-2HG production bears any functional significance or is due to promiscuous enzyme activity.

In eukaryotes, the degradation of the branch chain amino acids (BCAAs) valine and isoleucine yields propionyl-CoA, which can be converted to the short chain fatty acid (SCFA) propionate. In addition, propionate is produced by the gut microbiota during the breakdown of dietary fiber (Louis and Flint, 2017). Together with the other major SCFAs, acetate and butyrate, propionate forms an important source of energy for muscle, colon and liver (Reszko et al., 2003; Wong et al., 2006). Thus, propionate serves an important metabolic and physiological function. However, propionate is toxic when it accumulates to high levels in the blood, which occurs in patients with propionic acidemia that carry mutations in either of the two propionyl-CoA carboxylase subunits (Shchelochkov et al., 2012). These enzymes function together in the canonical, vitamin B12-dependent propionyl-CoA breakdown pathway that leads to the production of succinyl-CoA, which can anaplerotically enter the TCA cycle to produce energy (Martini et al., 2003).

Ketone bodies provide another important energy source under conditions where glucose is limiting, such as in diabetic patients, or on low carbohydrate, or “keto” diets. Ketone bodies are produced in the liver and are essential for energy metabolism in peripheral tissues such as muscle and the brain. There are three ketone bodies, two of which, acetoacetate and 3-hydroxybutyrate (3HB), can serve as energy sources, with 3HB being the most prevalent. Ketone bodies are produced from the breakdown of fatty acids and amino acids. Two amino acids, lysine and leucine, are exclusively ketogenic: degradation of leucine yields acetyl-CoA and acetoacetate, while breakdown of lysine yields 3HB. Furthermore, acetyl-CoA produced by leucine (and fatty acids) is an important precursor for ketone body production.

The nematode *C. elegans* has been a powerful model organism for decades, and research on this ‘simple’ animal has yielded great insights into development, aging, and other processes. Recently, *C. elegans* has also become a major model to study metabolism. Of its 20,000 or so protein-coding genes, more than 2,000 are predicted to encode metabolic enzymes. Of these, 1,314 have been annotated to specific metabolic reactions and incorporated into a genome-scale metabolic network model that can be used with flux balance analysis (FBA) to computationally model the animal’s metabolism (Yilmaz et al., 2020; Yilmaz and Walhout, 2016). Like in humans, vitamin B12 plays an important metabolic role in *C. elegans* propionate breakdown (MacNeil et al., 2013; Watson et al., 2013; Watson et al., 2014; Watson et al., 2016). Vitamin B12 is exclusively synthesized by bacteria and, perhaps, some archaea, and is therefore mostly acquired by diet. On bacterial diets low in vitamin B12, such as the standard laboratory diet of *Escherichia coli* OP50, *C. elegans* transcriptionally activates five genes that comprise an alternative propionate breakdown pathway, or propionate shunt (Bulcha et al., 2019; Giese et al., 2020; Watson et al., 2016). This shunt is activated only when high levels of propionate persist and detoxifies this SCFA to acetyl-CoA in the mitochondria (Bulcha et al., 2019).

Here, we use loss-of-function mutation in *C. elegans* D-2-hydroxyglutarate dehydrogenase, which converts D-2HG to αKG, to study endogenous D-2HG metabolism. We identified the *C. elegans* gene F54D5.12 as a one-to-one ortholog of human D2HGDH and named this gene *dhgd-1* for <u>D</u>-2-<u>h</u>ydroxy<u>g</u>lutarate <u>d</u>ehydrogenase. We find that D-2HG is produced by the propionate shunt, in the step in which HPHD-1 oxidizes 3-hydroxypropionate (3HP) to malonic semialdehyde (MSA). *dhgd-1* deletion mutants are embryonic lethal and have mitochondrial defects. Surprisingly, however, while mitochondrial defects can be explained by 3HP accumulation (Zhou et al., 2022), lethality is neither simply caused by accumulation of 3HP nor D-2HG. Therefore, loss of viability and mitochondrial defects are caused by distinct mechanisms and can be uncoupled. We find that *dhgd-1* mutant animals upregulate ketone body metabolism genes, suggesting that ketone bodies are limiting in these animals. Indeed, our metabolomic analysis shows that the breakdown of the ketogenic amino acids leucine and lysine is impaired in these mutants. Moreover, the ketone body 3HB can partially rescue *dhgd-1* mutant lethality. Altogether, our findings support a model in which ketone bodies are important for *C. elegans* viability.

## RESULTS

### DHGD-1 is a D-2-Hydroxyglutarate dehydrogenase

We first asked whether DHGD-1 is indeed a D-2HG dehydrogenase (**Figure 1A**). We obtained *dhgd-1(tm6671)* mutant animals that carry a large deletion in the 5’ end of the gene, which is predicted to result in a loss-of-function (**Figure 1B**). These animals are hereafter referred to as Δ*dhgd-1* mutants. By gas chromatography-mass spectrometry (GC-MS), we found that Δ*dhgd-1* mutants accumulate about five-fold more 2HG than wild type animals (**Figure 1C**). Human D2HGDH specifically oxidizes the D-form of 2HG to αKG (Yang et al., 2021). To test the stereo-specificity of *C. elegans* DHGD-1, we used chiral derivatization that allows the chromatographic separation of the D- and L-2HG enantiomers. While in wild type animals the ratio between L- and D-2HG is about one to one, Δ*dhgd-1* mutant animals predominantly accumulate the D-2HG enantiomer, confirming that DHGD-1 is indeed a functional ortholog of human D2HGDH (**Figure 1D**).

**Figure 1.**
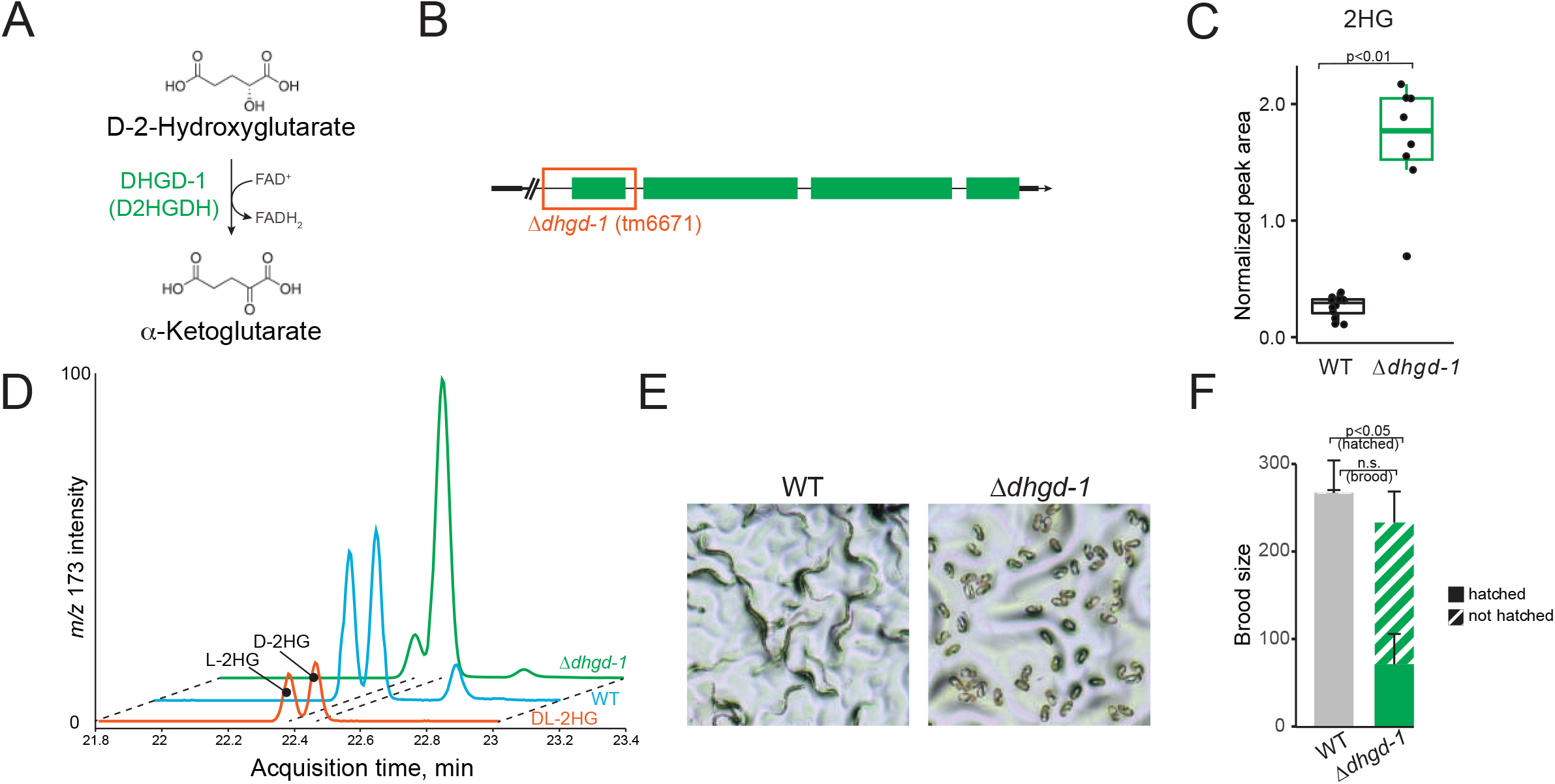
*Dhgd-1* encodes a D-2-hydroxyglutarate (D-2HG) dehydrogenase. (A) Metabolic reaction catalyzed by D-2HG dehydrogenase. (B) Schematic of deletion in *dhgd-1*(*tm6671*) mutants, referred to as Δ*dhgd-1* mutants. (C) GC-MS measurement of 2HG accumulation in Δ*dhgd-1* mutants and wild type (WT) animals. Each dot represents an independent biological replicate. (D) Discrimination between D- and L-2HG enantiomers by chiral GC-MS derivatization in WT and Δ*dhgd-1* mutant animals. Raw data are not normalized and therefore do not reflect 2HG concentration in different samples. (E,F) Embryonic lethality and brood size in Δ*dhgd-1* mutants. Brightfield images (E) and hatching (F). Scale bar 50 um. Bars in (F) represent mean and standard deviation of n=3 biological replicates. n.s. – not significant.

We found that Δ*dhgd-1* mutant animals have severe embryonic lethality as approximately two thirds of the embryos failed to hatch (**Figures 1E, F**). In addition, Δ*dhgd-1* mutant animals exhibit mitochondrial defects as their mitochondrial network is more fragmented and individual mitochondria are more rounded, which agrees with recently reported data (Zhou et al., 2022) (**Figure S1A**). These observations allow us to investigate the connection between organismal physiology and endogenous D-2HG metabolism.

### DHGD-1 functions in the propionate shunt

The *Δdhgd-1* mutant metabolome exhibits several additional changes in metabolite abundance, including lower accumulation of glutamate, αKG, succinate, and the lysine breakdown products 2-aminoadipate and glutarate, as well as higher abundance of lysine, β-alanine, 3-aminoisobutyrate, and the leucine breakdown product 3,3-hydroxymethylbutyrate (HMB) (**Figure 2A**). Importantly, 3HP, a metabolite unique to the propionate shunt (Watson et al., 2016), exhibited the greatest increase in abundance in Δ*dhgd-1* mutant animals, which suggests that loss of *dhgd-1* interferes with the function of the propionate shunt.

**Figure 2.**
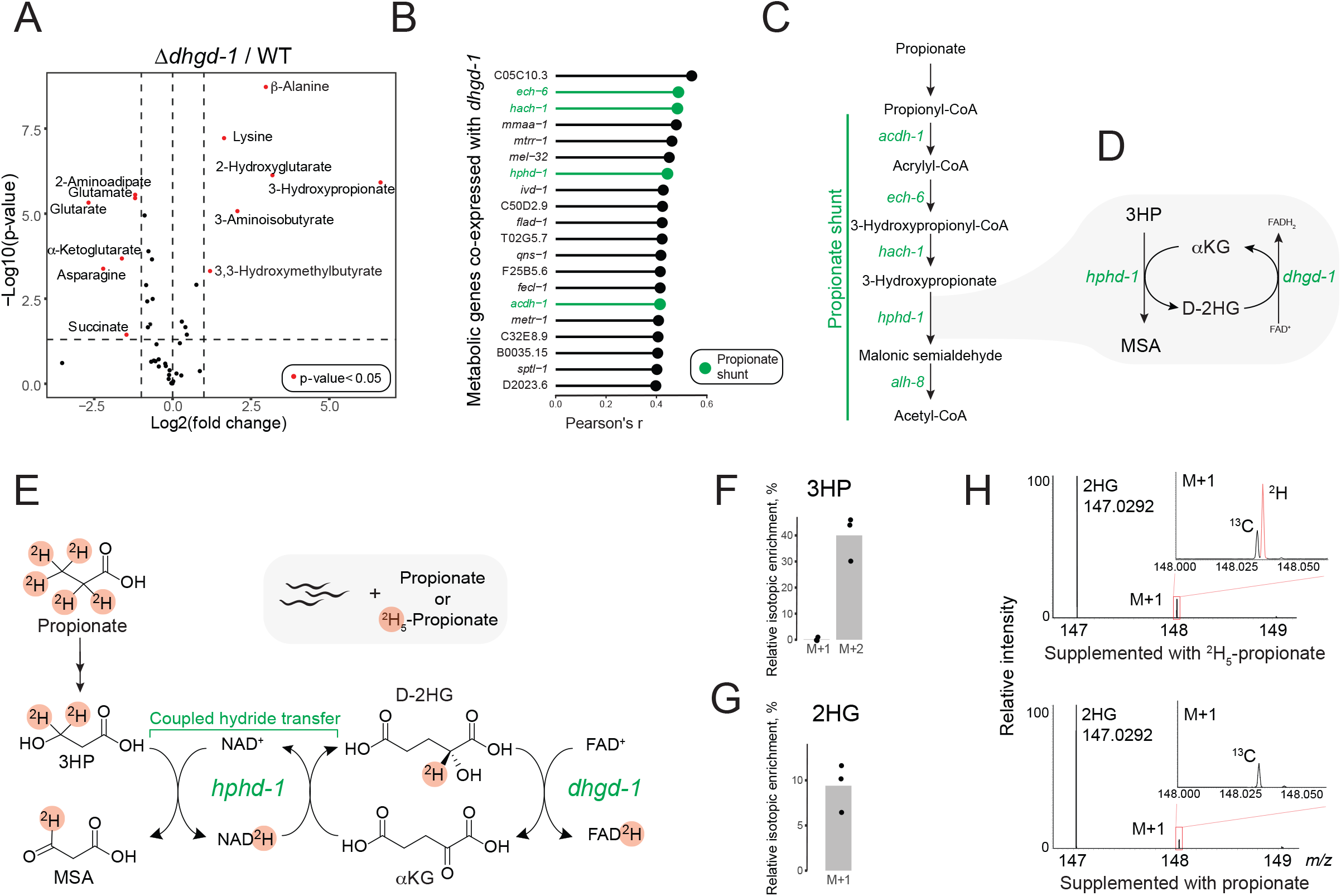
*Dhgd-1* functions in the propionate shunt pathway. (A) GC-MS profiling of metabolic changes in Δ*dhgd-1* mutants compared to WT *C. elegans*. P-values are Benjamini-Hochberg adjusted. (B) Metabolic genes most highly coexpressed with *dhgd-1*. Propionate shunt pathway genes are enriched with a false discovery rate (FDR) of 0.042 (Extended Data Table 1). (C,D) Proposed mechanism of D-2HG production (by HPHD-1) and recycling (by DHGD-1) during propionate degradation via the propionate shunt pathway. (E), Isotope tracing experiment with deuterium (^2^H)-labeled propionate and proposed reaction mechanism. HPHD-1 uses NAD^+^/NADH as a hydride shuttle in a coupled reaction yielding one equivalent each of MSA and D-2HG. (F,G) Fractional enrichment of 3HP (F) and 2HG (G) isotopologues. Bars indicate mean of n=3 biological replicates. (H) High performance liquid chromatography mass spectrometry (HPLC-MS) analysis of the M+1 isotope cluster in animals fed ^2^H_5_-propionate revealed robust incorporation of a single deuterium atom in 2HG, whereas the M+1 in animals fed propionate was exclusively from natural abundance of ^13^C.

By using a compendium of publicly available *C. elegans* expression profiles, we found that four of the five propionate shunt genes are strongly coexpressed with *dhgd-1* in a gene expression compendium with data from multiple growth conditions and genetic backgrounds (**Figures 2B,C, Table S1**)(Reece-Hoyes et al., 2013). One of these, *hphd-1*, is an ortholog of human ADHFE1, which produces D-2HG in a reaction coupled to oxidation of the neurotransmitter and psychoactive drug γ-hydroxybutyrate (Struys et al., 2005). Therefore, we hypothesized that HPHD-1 reduces αKG to D-2HG when it oxidizes 3HP to MSA (Watson et al., 2016), and that DHGD-1 recycles D-2HG back to αKG (**Figure 2D**). Since HPHD-1 harbors a highly conserved Rossman fold that can bind nucleic acid cofactors such as NAD+, we predicted that HPHD-1 uses NAD+/NADH to shuttle a hydride from 3HP to D-2HG (**Figure 2E**) (Gaona-Lopez et al., 2016). If true, this leads to the prediction that D-2HG is derived from propionate degradation. To directly test this, we performed a stable isotope tracing experiment in which animals were supplemented with either propionate or deuterated ^2^H_5_-propionate. Supplementing ^2^H_5_-propionate produced both deuterated 3HP and D-2HG, while the bacterial diet alone did not contain any detectable 3HP or 2HG, showing that these conversions happen in the animal (**Figures 2F-H** and **Figure S2**). This result demonstrates that production of D-2HG is coupled to oxidation of 3HP in the propionate shunt. We therefore conclude that DHGD-1 functions in the propionate shunt, and that its dysfunction leads to impaired flux through this pathway, resulting in accumulation of shunt intermediates 3HP and 2HG, as well as mitochondrial dysfunction and embryonic lethality.

### *hphd-1* RNAi and vitamin B12 supplementation rescue embryonic lethality of Δ*dhgd-1* mutants

High levels of 3HP cause mitochondrial defects in Δ*dhgd-1* mutants (Zhou et al., 2022). Do these defects cause embryonic lethality, or could this be caused by accumulation of 2HG? To test this, we knocked down *hphd-1*, which we predicted to reduce 2HG but not 3HP levels (**Figure 3A**). Indeed, RNAi of *hphd-1* reduced 2HG to levels that are similar to wild type animals but did not affect 3HP levels (**Figure 3B**). Remarkably, RNAi of *hphd-1* rescued lethality of *dhgd-1* mutants (**Figure 3C**). This result shows that embryonic lethality correlates with 2HG accumulation, and that this phenotype can be uncoupled from mitochondrial defects that are not rescued by *hphd-1* perturbation (Zhou et al., 2022). Both *dhgd-1* mutation and *hphd-1* RNAi block HPHD-1 function (Zhou et al., 2022). Because *hphd-1* RNAi does not cause embryonic lethality, we conclude that lack of flux through the propionate shunt is not sufficient to cause embryonic lethality.

**Figure 3.**
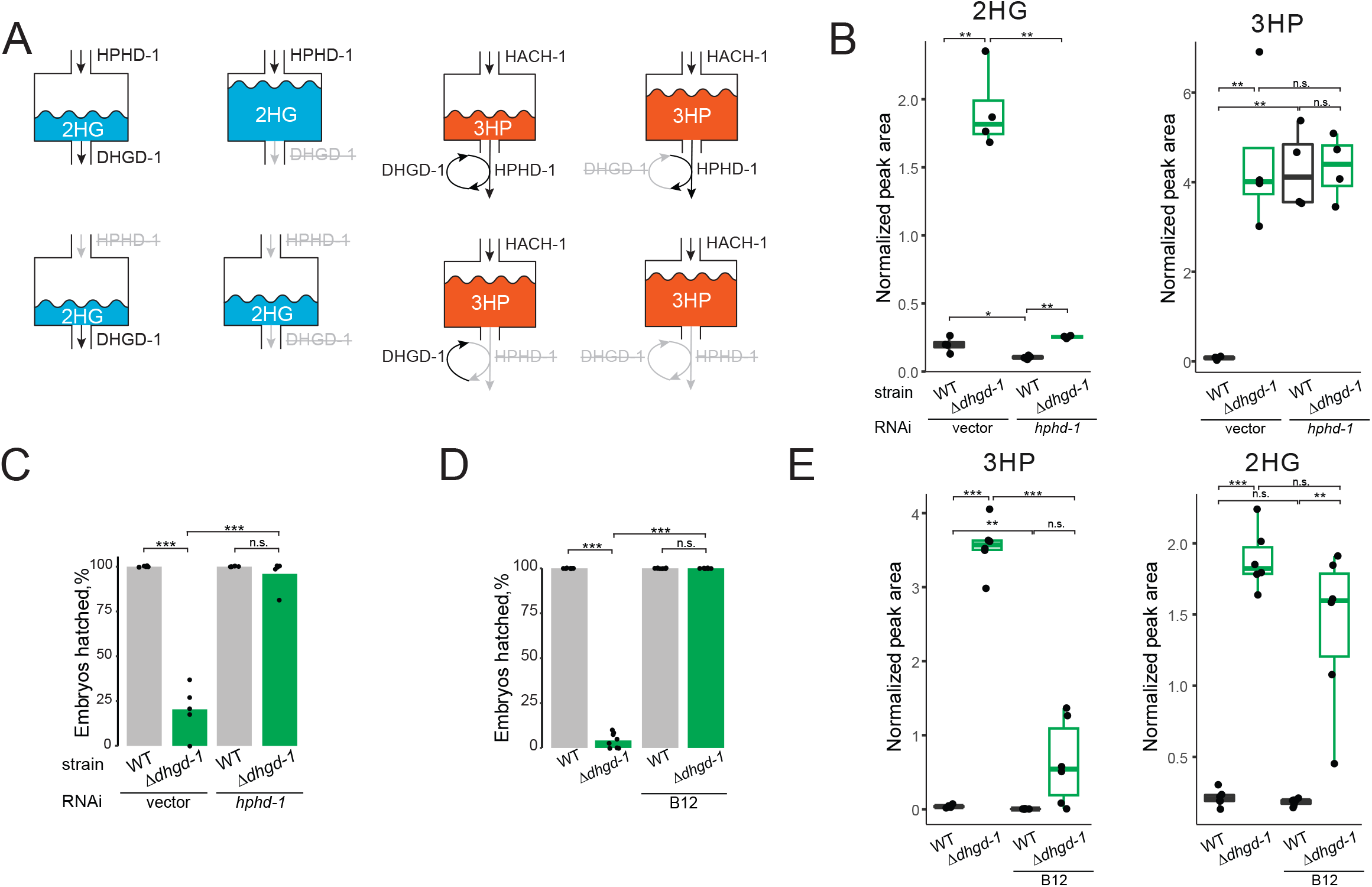
Rescue of lethality in Δ*dhgd-1* mutants by vitamin B12 supplementation and *hpdh-1* RNAi. (A) Schematic of HPHD-1 and DHGD-1 contributions to 2HG (blue) and 3HP (orange) accumulation. HPHD-1 is a main source of 2HG, and therefore its knockdown prevents 2HG accumulation in case of DHGD-1 dysfunction. 3HP is expected to accumulate if either DHGD-1 or HPHD-1 are perturbed because these reactions are coupled to facilitate 3HP oxidation. (B) 2HG and 3HP abundance in Δ*dhgd-1* mutants upon RNAi of *hphd-1*. Boxplot midline represents median of four independent biological replicates (dots). (C) *hphd-1* RNAi rescues lethality in Δ*dhgd-1* mutants. The RNAi-compatible *E. coli* OP50 strain(Xiao et al., 2015) was used because conventional RNAi-compatible *E. coli* HT115 rescued embryonic lethality of Δ*dhgd-1* animals. Each dot represents an independent biological replicate and bars indicate means. (D) Vitamin B12 rescues lethality in Δ*dhgd-1* mutants. Each dot represents an independent biological replicate and bars indicate means. (E) 3HP and 2HG abundance in Δ*dhgd-1* mutants supplemented with vitamin B12. Boxplot midline represents median of independent biological replicates (dots). All panels: *p<0.05, **p<0.01, ***p<0.001 (unpaired t-test).

Flux through the propionate shunt is transcriptionally repressed by vitamin B12, which enables flux through the canonical propionate degradation pathway (Bulcha et al., 2019; Watson et al., 2014; Watson et al., 2016). We found that supplementation of vitamin B12 rescued both mitochondrial defects and embryonic lethality in Δ*dhgd-1* mutants (**Figure 3D, Figure S1B**). Surprisingly, while, as expected, 3HP levels went down in Δ*dhgd-1* mutants upon supplementation of vitamin B12, 2HG levels remained high (**Figure 3E**). This observation suggests that there is still some flux through the propionate shunt pathway in Δ*dhgd-1* mutant animals supplemented with vitamin B12. More importantly, this result indicates that high levels of 2HG are not sufficient to elicit embryonic lethality in these mutants. Therefore, we hypothesized that vitamin B12 supplementation may circumvent detrimental effects of 2HG accumulation.

### Loss of *dhgd-1* may cause lethality by impairing ketone body production

To gain insight into the mechanism by which loss of *dhgd-1* causes embryonic lethality and how this could be rescued by vitamin B12 supplementation, we used expression profiling by RNA sequencing (RNA-seq). Overall, 315 and 183 genes are induced and repressed by loss of *dhgd-1*, respectively (**Table S2**). WormFlux (Yilmaz and Walhout, 2016) pathway enrichment analysis using pathways defined by WormPaths (Walker et al., 2021) revealed two main insights. First, the propionate shunt is not fully repressed by vitamin B12 in Δ*dhgd-1* mutants (**Figures 4A, B, Table S3**). This observation confirms that the propionate shunt remains active in the presence of vitamin B12 and can explain why 2HG levels remain high in these animals (**Figure 3E**). Second, ketone body metabolism genes are upregulated in Δ*dhgd-1* mutants with or without vitamin B12 compared to wild type and are downregulated in both Δ*dhgd-1* mutants and wild type by vitamin B12 in relation to their respective genotypes (**Figures 4A, B**). Therefore, we hypothesized that perturbed ketone body metabolism may explain embryonic lethality of Δ*dhgd-1* mutant animals. This hypothesis is supported by our metabolomic analysis, which found differential accumulation of metabolites in the breakdown of the ketogenic amino acids, leucine and lysine, in Δ*dhgd-1* mutants: lysine levels are increased while its breakdown products 2-aminoadipate and glutarate are decreased, and the leucine breakdown product HMB is increased (**Figures 2A** and **S3**). This therefore suggests that the breakdown of ketogenic amino acids lysine and leucine in Δ*dhgd-1* mutants is impaired at initial and intermediate steps, respectively, resulting in lower levels of ketone bodies. To directly test this hypothesis, we supplemented Δ*dhgd-1* mutants with the ketone body 3HB and found that it partially rescued embryonic lethality (**Figure 4C**). This result supports the interpretation that impaired ketone body metabolism (**Figure 4D**) causes embryonic lethality in Δ*dhgd-1* mutant animals.

**Figure 4.**
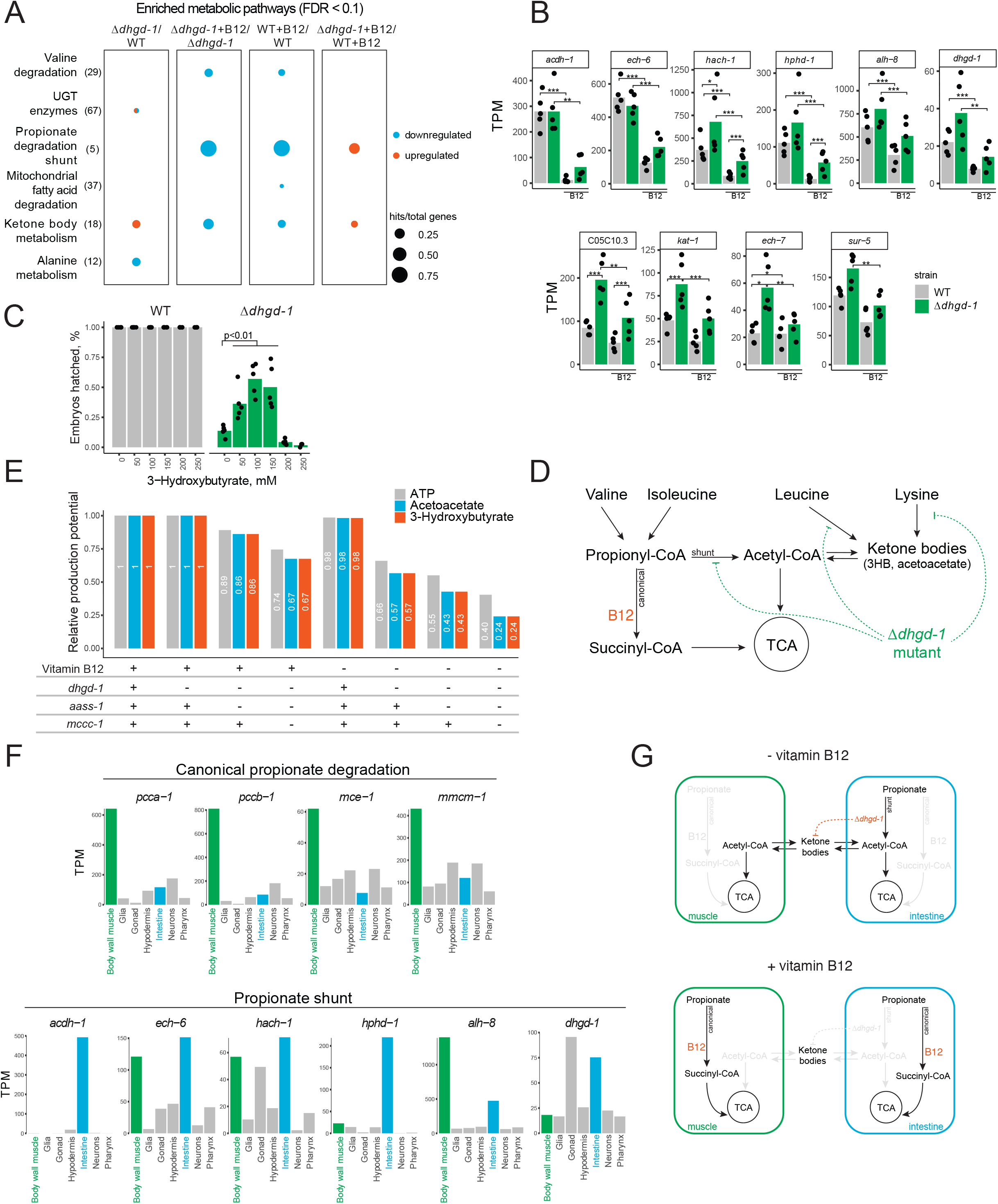
Loss of *dhgd-1* activates the expression of ketone body metabolism genes. (A) Gene expression pathway enrichment analysis for animals with and without vitamin B12 (B12). (B) RNA-seq data for propionate shunt (top) and ketone body metabolism genes (bottom). TPM – transcripts per million. (C) 3HB supplementation partially rescues lethality of Δ*dhgd-1* mutant animals. (E) Model for the effect of *dhgd-1* mutation on ketone body production. (F) Contribution of propionate shunt (*dhgd-1*), lysine (*aass-1*) and leucine (*mccc-1*) degradation pathways and vitamin B12 to the generation of energy (ATP) and ketone bodies (3HB and acetoacetate). Metabolite production potentials were estimated by maximizing their output fluxes in corresponding FBA formulations. (G) Tissue expression of genes comprising the canonical vitamin B12-dependent propionate degradation pathway and propionate shunt pathway from a published single cell RNA-seq dataset (Cao et al., 2017). (G) Model for ketone body exchange between *C. elegans* intestine and muscle in the presence and absence of vitamin B12.

Why are ketone bodies required for viability in Δ*dhgd-1* mutant animals? In mammals, ketone bodies are important carriers of energy (Fukao et al., 2014). Therefore, we hypothesized that *dhgd-1* may be required for energy production. Since lysine and leucine breakdown are impaired in Δ*dhgd-1* mutants, we modeled the impact of the combined loss of *dhgd-1* function with impaired lysine and leucine degradation on the production of both ketone bodies and energy. Specifically, we used FBA with the genome-scale *C. elegans* metabolic network model iCEL1314, and used maximization of ketone body or energy production as an objective function (Yilmaz et al., 2020; Yilmaz and Walhout, 2016). FBA predicted that energy and ketone body production are not affected by lack of vitamin B12 (loss of canonical propionate degradation pathway) or *dhgd-1* mutation (loss of the shunt pathway) alone (**Figure 3E**). However, on a diet low in vitamin B12, FBA predicted that loss of *dhgd-1* reduced both ketone body and energy production (**Figure 4E**). This result indicates that two propionate breakdown pathways may be important sources of energy. Indeed, the canonical propionate pathway anaplerotically replenishes TCA cycle by providing succinyl-CoA, while the product of the shunt, acetyl-CoA, is another TCA cycle intermediate and key ketone body precursor (Shi and Tu, 2015) (**Figure 4D**). Importantly, the modeling predicted that the additional loss of lysine and leucine degradation exacerbated the reduction in ketone body and energy production (**Figure 4E**). This observation provides further support for our hypothesis that ketone body production is impaired in Δ*dhgd-1* mutant animals (**Figure 4D**). Additionally, FBA modeling indicates that ketone bodies are an important source of energy in *C. elegans* on a standard laboratory diet low in vitamin B12, and that the propionate shunt plays a key role in ketone body production.

### A model for propionate catabolism in ketone body production and use in *C. elegans*

What is the physiological role of ketone body metabolism in *C. elegans*? In humans, ketone bodies are synthesized by the liver by the breakdown of fatty acids and the ketogenic amino acids lysine and leucine. The ketone body precursor, acetyl-CoA is converted into ketone bodies that are transported to tissues with high energetic demands (brain and muscle) (Puchalska and Crawford, 2017). Therefore, we hypothesized that, as in humans, ketone bodies may provide an energy source for body wall muscle in *C. elegans* to support development.

We obtained insight into the potential function of ketone bodies in *C. elegans* by examining tissue-level RNA-seq data of early larvae (Cao et al., 2017) and embryos (Warner et al., 2019). A key observation was that the canonical propionate breakdown pathway genes are extremely highly expressed in body wall muscle, relative to other tissues (**Figure 4F and S4**). How does this observation relate to ketone body metabolism? A major function of muscle metabolism is to produce energy for movement, which is critical for the developing *C. elegans* embryos. The observation that the canonical propionate breakdown pathway is most highly expressed in body wall muscle therefore indicates that this SCFA may serve as an important energy source for this tissue by generating succinyl-CoA that can enter the TCA cycle to support ATP production. How does muscle tissue generate energy on diets low in vitamin B12, when flux through this canonical pathway is impaired? We know that low vitamin B12 activates the expression of the propionate shunt (Bulcha et al., 2019; Watson et al., 2016). However, inspection of the tissue-level RNA-seq data mentioned above revealed that the first shunt gene, *acdh-1*, is not expressed in body wall muscle and, therefore, that this pathway is not active in this tissue (**Figure4G and S4**). Instead, this gene is most highly expressed in the animal’s intestine (**Figure 4F and S4**) and, at lower levels, in the hypodermis (**Figure 4F**). In *C. elegans*, intestine and hypodermis function as both gut and liver, which are the primary sites of nutrient absorption, digestion, and metabolism (Kaletsky et al., 2018; Yilmaz et al., 2020). The production of propionyl-CoA by BCAA breakdown is likely not affected by vitamin B12 and therefore, likely continues to provide an important energy source to the animal, even on diets low in this cofactor. Therefore, we hypothesized that, on low-vitamin B12 diets, the propionate shunt produces acetyl-CoA in the intestine (and to a lesser extent hypodermis) and that this acetyl-CoA is converted to ketone bodies (**Figure 4H**). These ketone bodies can then diffuse to the muscle, where they are converted back to acetyl-CoA and oxidized via the TCA cycle to produce energy. It is not feasible to directly test this hypothesis in *C. elegans* because different tissues cannot be isolated for metabolomic analyses. Therefore, we analyzed the production and consumption potential of ketone bodies, which we previously calculated using flux potential analysis (Yilmaz et al., 2020), and found that body wall muscle is the greatest consumer of ketone bodies.

## DISCUSSION

In this study, we identified a metabolic and physiological function of D-2HG production and recycling in *C. elegans*. This recycling is important for the animal because it facilitates flux through the propionate shunt, and deletion in *dhgd-1*, which encodes the enzyme that converts D-2HG back to αKG, results in mitochondrial defects and embryonic lethality. However, these phenotypes are caused by different metabolic changes: mitochondrial dysfunction but not embryonic lethality is caused by high levels of 3HP (Zhou et al., 2022). We further discovered that ketone body metabolism is impaired in Δ*dhgd-1* mutants. We put forth a model in which both propionate and ketone bodies provide an important source of energy for developing *C. elegans*. Propionate (or propionyl-CoA) is produced by the breakdown of the BCAAs valine and isoleucine, and, potentially, odd-chain fatty acids, while ketone bodies are generated from the breakdown of leucine, lysine and even-chain fatty acids, which generate the ketone body precursor acetyl-CoA. Our data suggest that propionate may provide a source of acetyl-CoA that enables ketone body production by the propionate shunt on diets low in vitamin B12.

There are three different mechanisms by which embryonic lethality in Δ*dhgd-1* mutant animals can be rescued, and each of these can be explained by our model. First, vitamin B12 supplementation fully rescues lethality, and we propose that this is because vitamin B12 facilitates production of propionate-derived succinyl-CoA in the muscle. Succinyl-CoA can enter the TCA cycle to produce energy, thereby circumventing the need for ketone body production either by the shunt or by breakdown of lysine or leucine. Second, direct supplementation of the ketone body 3HB partially rescues, likely because it can diffuse to the muscle where it can be converted to acetyl-CoA to produce energy. Finally, RNAi of *hphd-1* also fully rescues lethality in Δ*dhgd-1* mutant animals. This is a key result showing that loss of *dhgd-1* does not cause lethality solely by impairing flux through the propionate shunt, because *hphd-1* functions in the same pathway, even in the same reaction coupled with *dhgd-1*. Our metabolomic data showed that loss of *dhgd-1* may impair lysine and leucine breakdown, which may occur as a result of 2HG accumulation, which is known to inhibit a variety of metabolic enzymes, including BCAT, a key enzyme in the breakdown of all three BCAAs (McBrayer et al., 2018). Altogether, our results indicate that *C. elegans* may rely on amino acids and propionate as a source of energy, and that on diets low in vitamin B12, the propionate shunt is an important producer of the ketone body precursor, acetyl-CoA. We have previously shown that the propionate shunt plays an important role in the detoxification of propionate, high levels of which are detrimental to *C. elegans*, as they are in humans (Bulcha et al., 2019; Watson et al., 2016). Therefore, the results presented here indicate that the propionate shunt serves an important dual purpose: the detoxification of excess propionate, and the production of acetyl-CoA and energy.

## Acknowledgements

We thank members of the Walhout lab, especially Yong-Uk Lee, for discussion and critical feedback on the manuscript.

## Author contributions

O.P. and A.J.M.W. conceived the project. O.P. performed all experiments and analyses with help from H.Z., G.G and X.L. (RNA-seq), S.N. (gene coexpression analysis), T.L. (3-HB supplementation) and B.F. (HPLC-MS). S.L.Y. performed flux balance analysis. O.P. and A.J.M.W. wrote the paper with input from all other authors.

## STAR METHODS

### RESOURCE AVAILABILITY

#### Lead contact

Further information and requests for reagents may be directed to and will be fulfilled by the Lead Contact, A.J.M. Walhout.

#### Materials availability

*hphd-1* RNAi and vector control *E. coli* OP50 (xu363) strains are available upon request.

#### Data and code availability

Sequencing data have been deposited in GEO under accession code GSE201645. Any additional information required to reanalyze the data reported in this paper is available from the lead contact upon request.

### EXPERIMENTAL MODEL AND SUBJECT DETAILS

#### *C. elegans* and *E. coli* strains

For maintenance, *C. elegans* strains were grown on nematode growth medium (NGM) seeded with *E. coli* OP50 and supplemented with 64 nM adenosylcobalamin (vitamin B12). Experimental plates contained vitamin B12 if specified. The mutant strain F54D5.12*(tm6671)* was provided by the National Bioresource Project for Nematode *C. elegans*(Mitani, 2009) and was backcrossed three times to N2. We named F54D5.12 *dhgd-1* for <u>D</u>-2-<u>H</u>ydroxy<u>g</u>lutarate <u>d</u>ehydrogenase. Wild type N2 *C. elegans* and bacterial strains *E. coli* OP50 and *E. coli* HT115 were obtained from the Caenorhabditis Genetics Center (CGC,) which is funded by NIH Office of Research Infrastructure Programs (P40 OD010440). The RNAi-compatible *E. coli* OP50(xu363) was provided by the Xu lab(Xiao et al., 2015).

## METHOD DETAILS

### Metabolite extractions

*C. elegans* cultures maintained on *E. coli* OP50 diet supplemented with 64 nM vitamin B12 were synchronized using sodium-hydroxide-buffered sodium hypochlorite treatment. L1 larvae were plated onto NGM agar plates for specified treatment (*e*.*g*., vitamin B12 supplementation, RNAi) and harvested as 1-day-old gravid adults. Animals used for experiments described in Figure 1C were cultured in liquid S-medium (Hibshman et al., 2021) in conditions described in the isotope tracing section of the methods. First day gravid adults were washed three times with M9 buffer and 50 ul of the packed animal pellet was flash-frozen in a dry ice/ethanol bath and stored at -80°C. Next, samples were mixed with 1 mL 80% methanol and 0.5 mL of 200- to 300-μm acid-washed glass beads (MilliporeSigma) and homogenized using a FastPrep-24 bead beater (MP Biomedicals), with intermittent cooling in dry ice/ethanol bath. Samples were then extracted for 15 min, then centrifuged for 10 min at 20,000 × *g*, and the supernatant was used immediately or stored at -80°C.

### Targeted quantification of metabolites using gas chromatography-mass spectrometry (GC-MS)

250 ul of animal extracts were dried under vacuum using a SpeedVac concentrator SPD111V (Thermo Fisher Scientific). Derivatization of dried samples was performed by adding 20 ul pyridine and 50 μL *N-* methyl-*N*-(trimethylsilyl)trifluoroacetamide (MSTFA, MilliporeSigma) and incubating samples for three hours at 37° C, followed by five hours incubation at room temperature. In the experiments that also targeted αKG, dried samples received 20 μL of 20 mg/mL methoxyamine hydrochloride in pyridine (MilliporeSigma) and were incubated at 37° C for one hour before reaction with MSTFA derivatization. Measurements were performed on an Agilent 7890B single quadrupole mass spectrometry coupled to an Agilent 5977B gas chromatograph (GC-MS) with an HP-5MS Ultra Inert capillary column (30 m × 0.25 mm × 0.25 μm). The inlet temperature was set to 230°C, the transfer line was at 280°C, and the MS source and quadrupole were at 230°C and 150°C, respectively. The oven was programmed to start at 80°C, hold for 2 min, and ramp-up at 5°C/min until 280°C. Each metabolite was identified based on retention time, one quantifier, and two qualifier ions that were manually selected using a reference compound. Peak integration and quantification of peak areas was done using MassHunter software, blank subtraction and normalization to total quantified metabolites were done in R software.

### Relative quantification of D- and L-2HG

To distinguish the two enantiomers of 2HG we adapted a previously published protocol(Li and Tennessen, 2019). First, 300 ul of *C. elegans* metabolite extract was evaporated to dryness in glass inserts. 50 ul of R-(-)-butanol and five ul of 12N hydrochloric acid were added to each insert and incubated at 90°C with shaking for three hours. Samples were cooled to room temperature and transferred into glass tubes containing 400 ul hexane. After extraction 250 ul of organic phase were transferred into a new glass insert and evaporated to dryness. Next, 30 ul of pyridine and 30 ul of acetic anhydride were added to each sample and allowed to incubate for one hour at 80°C with shaking. Samples were dried, resuspended in 60 ul of hexane, and immediately analyzed by GC-MS using targeted method settings, with exception of oven ramp, which was run from 80 to 190°C at the rate of 5°C/min and then until 280°C at 15°C/min. 173 m/z ion was used for quantification of both D- and L-2HG.

### Brood size and hatching assays

Seven L4-stage animals per strain/condition were placed on individual 3.5 cm plates and transferred to a new plate every 24 hours until animals stopped laying eggs. Plates with embryos were incubated for 24 hours and then the number of hatched L1 larvae and unhatched embryos were counted and averaged for all animals. Brood counts for animals that died or crawled off the plate before egg-laying was complete were excluded. The experiment was repeated three times.

### Mitochondrial network imaging

Wild type and Δ*dhgd-1 C. elegans* strains were crossed to a strain with fluorescently labeled mitochondria and nuclei *Pmyo-3::GFP*mito; *Pmyo-3::lacZ::GFP*(nls) in body wall muscles. At least 10 L4 animals per condition were imaged using Nikon A1 point-scanning confocal microscope with 561 nm laser. Imaging was performed using an Apo TIRF, N.A. 1.49, 60x oil immersion objective in galvano imaging mode.

### Gene coexpression analysis

A ranked list of metabolic genes that are coexpressed with *dhgd-1* was extracted from a compendium of 169 expression datasets. Briefly, z-normalized expression datasets with at least 10 conditions were combined to form a global coexpression matrix. Correlation in expression between metabolic gene pairs across the compendium was calculated as Pearson Correlation Coefficient. Gene set enrichment analysis (GSEA) was performed on the ranked coexpression list using the PreRank module of GSEA (Subramanian et al., 2005) with pathway-to-gene annotations from WormPaths (Walker et al., 2021) and the significance cutoff set at a FDR of less than or equal to 0.05.

### Deuterium isotope tracing

Gravid adults were treated with sodium-hydroxide-buffered sodium hypochlorite solution to obtain synchronous L1 populations. Approximately 50,000 L1 animals were added per 50 ml Erlenmeyer flask containing 10 ml of K-medium with modified salt concentrations (51 mM NaCl, 32 mM KCl, 3 mM CaCl2, 3 mM MgSO4) and *E. coli* OP50 pellet from a 100 ml of overnight culture. Cultures were incubated at 20°C with shaking at 180 rpm. Sodium hydroxide-neutralized propionic acid (MilliporeSigma) or ^2^H_5_-propionic acid sodium salt (Cambridge Isotope Laboratories) were added to the cultures when animals reached the late L4/young adult stage to a final concentration of 20 mM. *C. elegans* were harvested at the gravid adult stage (about 12 h after supplementation), washed three times, flash-frozen in ethanol/dry ice bath, and stored at -80°C. Metabolite extraction, derivatization, and GC-MS methods were as described above. M+0 3HP and 2HG were quantified as *m/z* 219 and *m/z* 247, respectively. Relative isotopic enrichment was calculated as 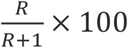, where R is a difference between abundances (normalized to the M+0 ion) of isotopologues in a sample with labeled propionate and a sample with unlabeled propionate supplement.

### HPLC-MS analysis

Reversed-phase chromatography was performed using a Vanquish HPLC system controlled by Chromeleon Software (ThermoFisher Scientific) and coupled to an Orbitrap Q-Exactive HF mass spectrometer controlled by Xcalibur software (ThermoFisher Scientific). Extracts prepared as described above were separated on a Thermo Scientific Hypersil Gold column (150 mm x 2.1 mm, particle size 1.9 µm) maintained at 40 °C with a flow rate of 0.5 mL/min. Solvent A: 0.1% formic acid in water; solvent B: 0.1% formic acid in acetonitrile. A/B gradient started at 1% B for 3 min after injection and increased linearly to 98% B at 20 min, followed by 5 min at 98% B, then back to 1% B over 0.1 min and finally held at 1% B for an additional 2.9 min to re-equilibrate the column. Mass spectrometer parameters: spray voltage (−3.0 kV, +3.5 kV), capillary temperature 380 °C, probe heater temperature 400 °C; sheath, auxiliary, and sweep gas 60, 20, and 2 AU, respectively. S-Lens RF level: 50, resolution 120,000 at m/z 200, AGC target 3E6. Samples were injected and analyzed in negative and positive electrospray ionization modes with m/z range 117-1000. Analysis was performed with Xcalibur QualBrowser v4.1.31.9 (Thermo Scientific).

### Metabolite supplementation

For experiments using vitamin B12 supplementation, NGM agar plates were supplemented with 64 nM adenosylcobalamin before pouring plates. DL-3-hydroxybutyric acid sodium salt (MilliporeSigma) was supplemented in indicated concentrations to NGM agar media.

### Embryonic lethality assays

Gravid adults were treated with sodium-hydroxide-buffered sodium hypochlorite solution, and released embryos were washed and incubated in M9 buffer for 18 hours to obtain synchronized L1 animals. Approximately 30 animals were placed onto 3.5 cm NGM plates (seeded with 100 ul of overnight bacterial culture one day before) and allowed to lay eggs. Adult animals were washed away with M9 buffer and approximately 200-300 embryos were transferred onto a new plate with the same supplements. After 24 hours hatched and not hatched animals were counted.

### RNAi assays

RNAi experiments were done using *E. coli* OP50(xu363) (Xiao et al., 2015) transformed with either empty vector L4440 or RNAi plasmid. Bacterial cultures were grown 18-20 hours and seeded onto NGM agar plates containing 2 mM isopropyl β-D-1-thiogalactopyranoside (IPTG), 50 μg/mL ampicillin, and used for metabolomics or phenotypic assays as described.

### Expression profiling by RNA-seq

200-300 synchronized gravid adults were harvested from *E. coli* OP50-seeded NGM plates with or without supplemented 64 nM adenosylcobalamin. Animals were washed three times with M9 buffer and total RNA from their bodies (excluding embryos) was extracted using the RNeasy kit (Qiagen), with an additional step of on-column DNase I (NEB) treatment. RNA quality was verified by agarose gel electrophoresis.

RNA-sequencing was performed as previously described (Giese et al., 2020). Briefly, multiplexed libraries were prepared using Cel-seq2 (Hashimshony et al., 2016). Two biological replicates were sequenced with a NextSeq 500/550 High Output Kit v2.5 on a Nextseq500 sequencer and three other replicates were sequenced on MGISEQ-2000. The libraries were first demultiplexed by a homemade python script, and adapter sequences were trimmed using trimmomatic-0.32 by recognizing polyA and barcode sequences. Then, the alignment to the reference genome was performed by STAR. Features were counted by ESAT (Derr et al., 2016) with pseudogenes discarded. The read counts for each gene were used in differential expression analysis by DESeq2 package in R 3.6.3(Love et al., 2014). Batch effects were corrected in DESeq2 statistical model by adjusting the design formula.

### Flux balance analysis

Flux balance analysis was done using established methods (Yilmaz and Walhout, 2016) with the *C. elegans* metabolic network model iCEL1314(Yilmaz et al., 2020), which can be accessed at http://wormflux.umassmed.edu/. Before simulations, the model was modified by associating HPHD-1 solely with the oxidation of 3HP(Watson et al., 2016). In all simulations, regular model constraints, such as allowed bacterial intake and forced growth-independent maintenance energy cost, were applied using reaction boundaries. With two additional constraints, glyoxylate shunt pathway was made irreversible and propionate secretion was prevented (i.e., by setting the lower boundary of reaction RM00479 and the upper boundary of reaction EX00163 as zero, respectively). A background metabolic activity was set using reaction BIO0107, which produces biomass with all possible macro-components(Yilmaz et al., 2020), as the objective function. The flux of this reaction was maximized first without any additional constraints.

Then, to obtain maximum biomass production potential at low B12 conditions, the flux of B12-dependent methionine synthase (MS) reaction (RC00946) was constrained to half of its value and the maximization was repeated. This constraint reduced the biomass flux (i.e., the maximum flux of BIO0107) by about 9% as expected (Yilmaz and Walhout, 2016). For subsequent simulations, the flux of BIO0107 was constrained to be at least at this level to represent the background metabolic activity, while the constraint on MS was removed (except for low B12 simulations, see below). FBA results were obtained using objective functions that maximized the fluxes of reactions RCC0005, EX00164, EX03197, which represent the generation of energy, and the production of acetoacetate and 3HB, respectively. For each objective, four FBA runs were first done using the following constraints: (i) no additional constraint was applied; (ii) DHGD-1 reaction (RM03534) was constrained to zero flux to represent the mutated *dhgd-1* and resulting removal of flux through the propionate shunt; (iii) *aass-1*, L-lysine-alpha-ketoglutarate reductase reaction (RM00716), was constrained to zero flux to represent the lack of lysine degradation; and (iv) methylcrotonyl-CoA carboxylase reaction (RM04138) was constrained to zero flux to represent the lack of leucine degradation. The constraint of each run was added in the respective order, while maintaining the constraints from previous runs. Finally, all four FBA runs were repeated with two additional constraints that represent low B12 conditions: RC00946 was constrained to half of its optimal value as described above and methylmalonyl-coA mutase reaction (RM00833) was constrained to zero value.

### QUANTIFICATION AND STATISTICAL ANALYSIS

All statistical details of experiments can be found in the figure legends.

## KEY RESOURCES TABLE

| REAGENT or RESOURCE | SOURCE | IDENTIFIER |
| --- | --- | --- |
| Bacterial and virus strains |  |  |
| *Escherichia coli* OP50 | CGC | N/A |
| *Escherichia coli* OP50 (xu363) | (Xiao et al., 2015) | N/A |
| Chemicals, peptides, and recombinant proteins |  |  |
| Coenzyme B12 (adenosyl cobalamin) | MilliporeSigma | Cat# C0884 |
| *N*-methyl-*N*-(trimethylsilyl)trifluoroacetamide (MSTFA) | MilliporeSigma | Cat#M-132 |
| Methoxyamine hydrochloride | MilliporeSigma | Cat# 226904 |
| DL-3-hydroxybutyric acid sodium salt | MilliporeSigma | Cat# H6501 |
| Propionic acid | MilliporeSigma | Cat#81910 |
| SODIUM PROPIONATE (D5, 98%) | Cambridge Isotope Laboratories | Cat#DLM-1601-1 |
| Deposited data |  |  |
| Transcriptome dataset (Effect of mutated *dhgd-1(tm6671)* on *C. elegans* metabolism) | This study | https://www.ncbi.nlm.nih.gov/geo/query/acc.cgi?acc=GSE201645 |
| Experimental models: Organisms/strains |  |  |
| *Caenorhabditis elegans* N2 (wild type) | CGC | N/A |
| *Caenorhabditis elegans* F54D5.12(*tm6671*) | (Mitani, 2009) | N/A |
| *Caenorhabditis elegans* Pmyo-3::*GFP*mito; Pmyo-3::*lacZ*::*GFP*(*nls*) | CGC | PD4251 |
| Software and algorithms |  |  |
| R: A Language for Data Analysis and Graphics, version 4.1 | http://www.r-project.org | N/A |
| WormFlux | http://wormflux.umassmed.edu/ | N/A |
| Other |  |  |
| Single quadrupole GC/MS | Agilent Technologies | 5977B GC/MSD |

## SUPPLEMENTAL FIGURE LEGENDS

**Figure S1.**
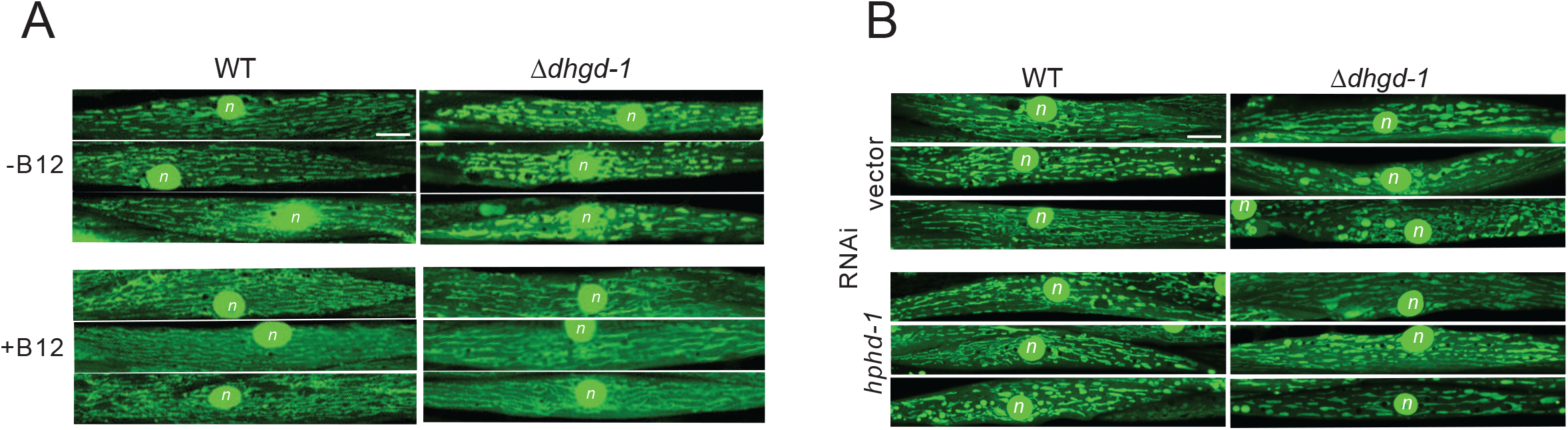
Mitochondrial defects in Δ*dhgd-1* mutants. (A,B) Representative images of mitochondria labelled with *Pmyo-3::GFP*mito in the wall body muscle of L4 larval stage animals. Defects in mitochondrial morphology of Δ*dhgd-1* mutant *C. elegans* is rescued by supplementing vitamin B12 (A) but not by *hphd-1* RNAi (B). Scale bar 10 um. Nuclei are marked with ‘n’.

**Figure S2.**
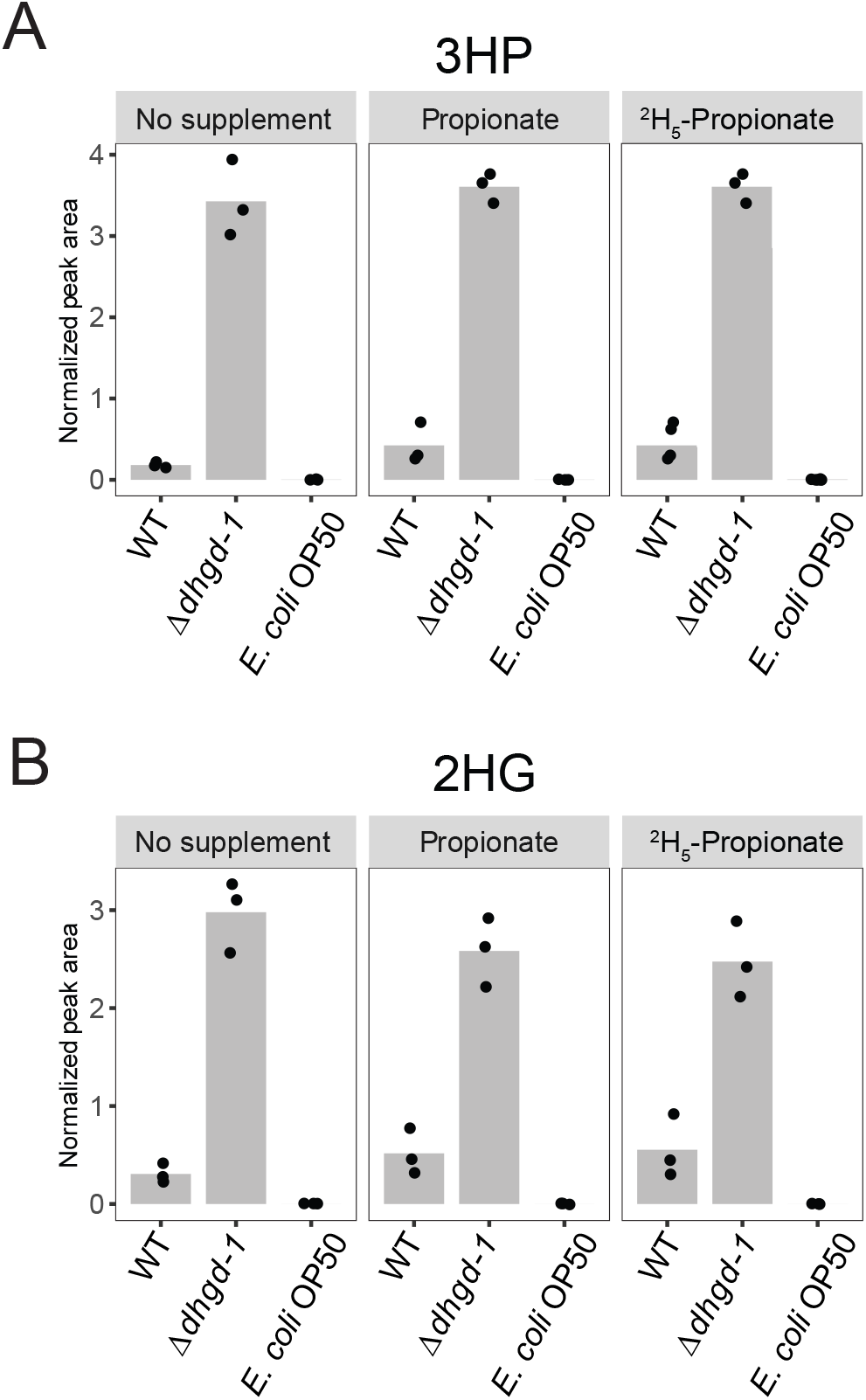
3HP and 2HG measurements by GC-MS. (A,B) GC-MS quantification of 3HP (A) and 2HG (B) in *C. elegans* and *E. coli* OP50 supplemented with propionate, ^2^H_5_-propionate or untreated. Bars represent mean, each dot represents an independent biological replicate,

**Figure S3.**
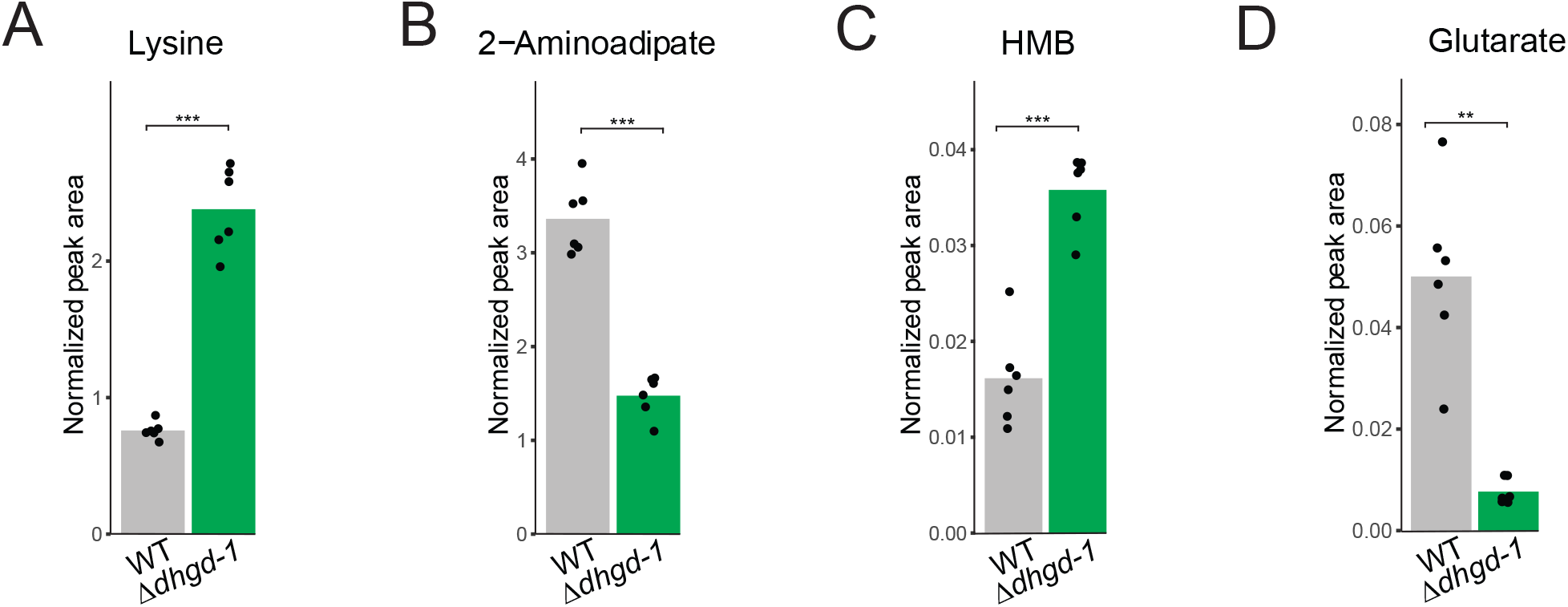
GC-MS measurement of intermediates in degradation of ketogenic amino acids lysine and leucine. (A,B,C,D) GC-MS quantification of lysine (A), 2-aminoadipate (B), HMB (C) and glutarate (D) in Δ*dhgd-1* mutants and wild type (WT) animals. Each dot represents an independent biological replicate, **p<0.01, ***p<0.001.

**Figure S4.**
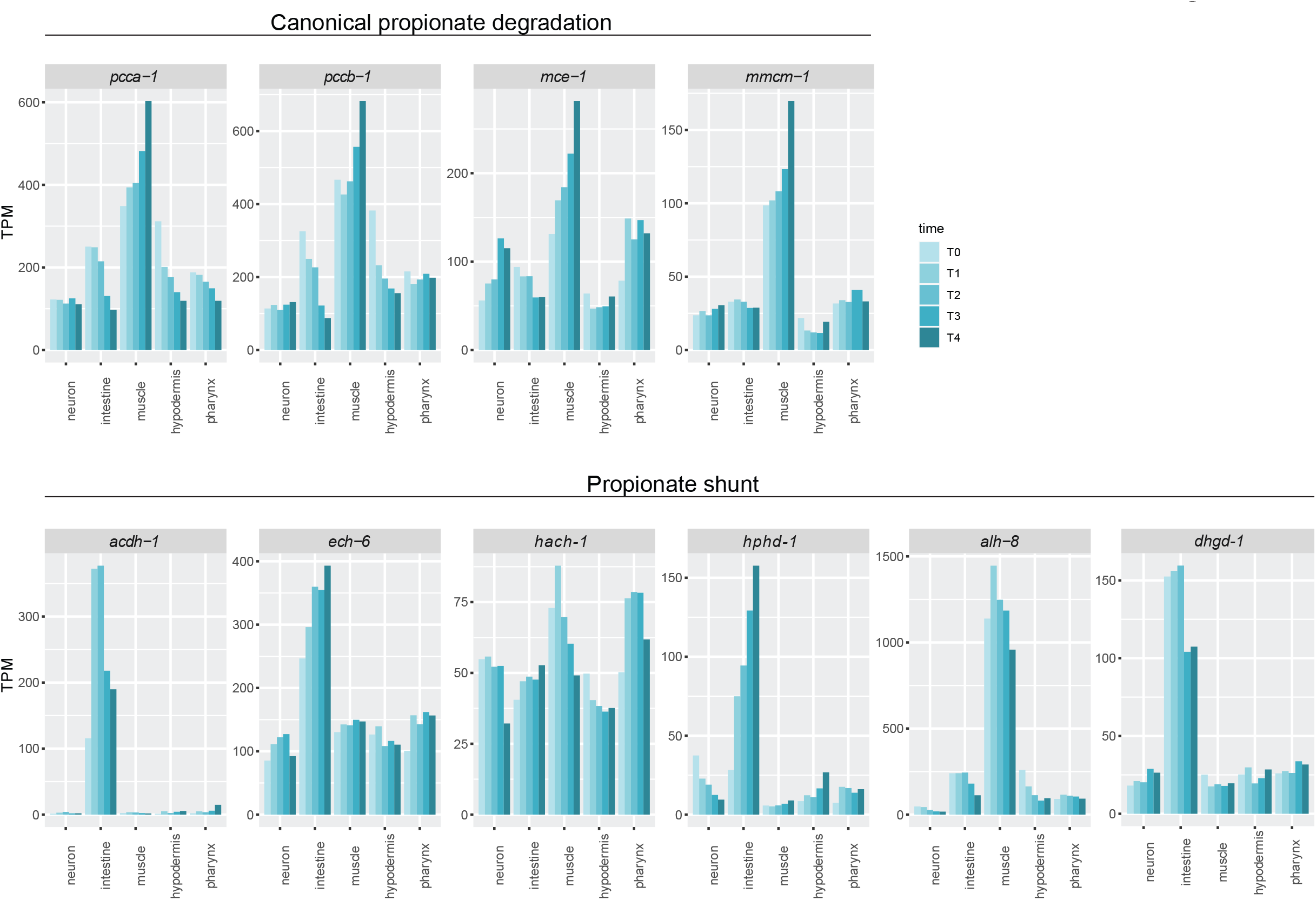
Embryonic tissue expression of genes comprising the two *C. elegans* propionate degradation pathways. Canonical, vitamin B12-dependent pathway (top) and propionate shunt (bottom) from a published dataset (Warner et al., 2019).

## SUPPLEMENTAL TABLES

**Table S1. Gene sets enriched in genes coexpressed with *dhgd-1***

**Table S2. Genes differentially expressed in Δ*dhgd-1* mutant *C. elegans* as compared to wild type**

**Table S3. WormFlux pathway enrichment analysis of genes differentially expressed in Δ*dhgd-1* mutant animals with and without vitamin B12**

